# How does Mg^2+^ modulate the RNA folding mechanism — a case study of G:C W:W Trans base pair

**DOI:** 10.1101/098079

**Authors:** Antarip Halder, Rohit Roy, Dhananjay Bhattacharyya, Abhijit Mitra

**Affiliations:** Center for Computational Natural Sciences and Bioinformatics (CCNSB), International, Institute of Information Technology (IIIT-H), Gachibowli, Hyderabad 500032, India; Computational Science Division, Saha Institute of Nuclear Physics (SINP), 1/AF,Bidhannagar, Kolkata 700064, India

## Abstract

Reverse Watson-Crick G:C base pairs (G:C W:W Trans) occur frequently in different functional RNAs. It is one of the few base pairs whose gas phase optimized isolated geometry is inconsistent with the corresponding experimental geometry. Several earlier studies indicate that accumulation of positive charge near N7 of guanine, through posttranscriptional modification, direct protonation or coordination with Mg^2+^, can stabilize the experimental geometry. Interestingly, recent studies reveal significant variation in the position of putatively bound Mg^2+^. This, in conjunction with recently raised doubts regarding some of the Mg^2+^ assignments near the imino nitrogen of guanine, is suggestive of the existence of multiple Mg^2+^ binding modes for this base pair. Our detailed investigation of Mg^2+^ bound G:C W:W Trans pairs, occurring in high resolution RNA crystal structures, show that they occur in 14 different contexts, 8 out of which display Mg^2+^ binding at the Hoogsteen edge of guanine. Further examination of occurrences in these 8 contexts led to the characterization of three different Mg^2+^ binding modes, (i) direct binding via N7 coordination, (ii) direct binding via O6 coordination and (iii) binding via hydrogen bonding interaction with the first shell water molecules. In the crystal structures, the latter two modes are associated with a buckled and propeller twisted geometry of the base pair. Interestingly, respective optimized geometries of these different Mg^2+^ binding modes (optimized at B3LYP) are consistent with their corresponding experimental geometries. Subsequent interaction energy calculations at MP2 level, and decomposition of its components, suggest that for G:C W:W Trans, Mg^2+^ binding can fine tune the base pair geometries without compromising with their stability. Our results, therefore, underline the importance of the mode of binding of Mg^2+^ ions in shaping RNA structure, folding and function.

## Introduction

Consequent to the discovery of their enzymatic roles,(*1, 2*) RNA has been found to be associated with numerous biophysical processes.(*3*) To execute these cellular processes, including regulation of gene expression and protein synthesis, RNA molecules are required to be folded in functionally competent structures which display a diverse repertoire of noncanonical base pairs.(*4*) The unique geometry and stability of these noncanonical base pairs shape up the characteristic features of different structural motifs in RNA.(*5, 6*) Hence, recognizing the role of different noncanonical base pairs is important for developing a comprehensive understanding of the sequence-structure-function relationship in RNA.

In this context, quantum mechanics based theoretical studies of the intrinsic properties of these base pairs have achieved a remarkable success.(*7, 8*) This was possible since in most of the cases, minimum energy geometries of the isolated base pairs obtained in gas phase are very much consistent with their geometry as observed within the crystal environment (experimental geometry).(*9*) Naturally, those noncanonical base pairs, whose minimum energy structures are significantly different from their experimental geometry, become the subject of interest. Identifying the physicochemical factors that stabilize these ‘away from equilibrium’ local geometries within the crystal environment constitute an important problem. On the one hand it provides an insight into RNA’s structural dynamics, and on the other, it provides a testbed for designing nucleic acid based nano devices with switching potential. One classical example of such a noncanonical base pair is the Levitt base pair,(*10*) the conserved interaction between G15 and C48 residues in tRNA. This crucial tertiary interaction is located at the elbow of the L-shaped structure of cytosolic tRNAs, and mediates the interaction between the D and V arms. Within the crystal structure, the G15-C48 interaction is found to be present in reverse Watson-Crick geometry (RWC), stabilized by 2 hydrogen bonds. However, on isolated geometry optimization in gas phase, it converges to a bifurcated geometry, stabilized by two bifurcated hydrogen bonds(*11*) (Figure 1).^1^

**Figure 1:**
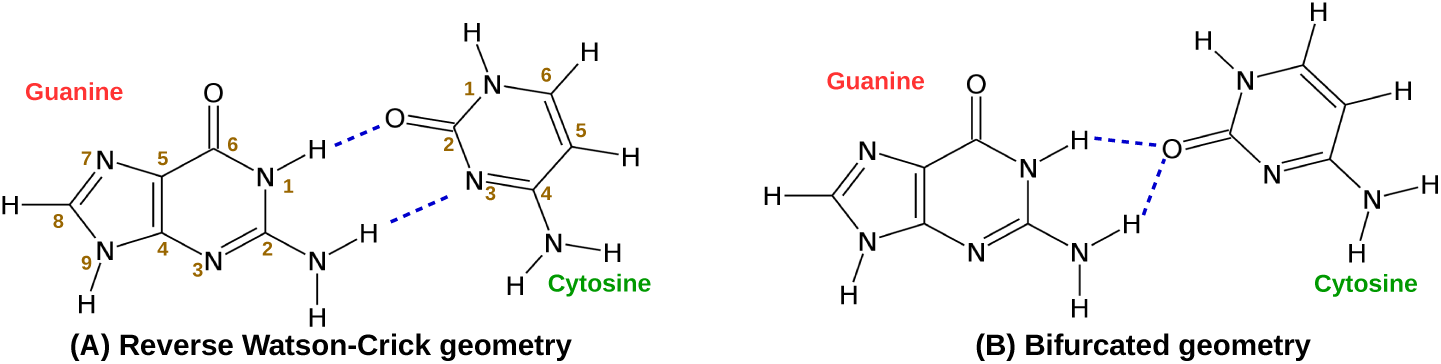
Schematic representation of the reverse Watson-Crick (RWC) and bifurcated (BF) geometry of the Levitt base pair. Hydrogen bonds are shown as blue broken lines.

Analyzing tRNA crystal structures, Oliva and co-workers have identified that positive charge build up at the hoogsteen edge of guanine (at N7 position), in the form of coordination with Mg^2+^ ion or posttranscriptional archaeosine modification, stabilizes the RWC geometry of Levitt base pair.(*13*) Binding of other divalent cations like Mn^2+^ and Co^2+^ at the N7 position of guanine also have similar effect on the RWC geometry of G:C W:W Trans pair.(*14*) However, occurrence of G:C W:W Trans base pair is not limited to G15-C48 of tRNA, rather they occur recurrently in 23S rRNAs.(*15*) They are also present in class II preQ1 riboswitches and participate in the recognition mechanism.(*16*) Chawla and co-workers have analyzed all such occurrences of G:C W:W Trans pair in high resolution RNA crystal structures and have found that, two third of the total occurrences of G:C W:W Trans are involved in higher order interactions (*e.g.*, base triples, base quartets, interaction with ordered surrounding water molecules and interaction with phosphate backbone) that stabilize its RWC geometry.(*15*) In our earlier work, we have shown that, direct protonation at the N7 position of guanine is sufficient to stabilize the RWC geometry.(*17*) Although protonation of nucleobases is thermodynamically unfavorable,^2^ the importance of charged nucleobases in modulating structure(*19–22*) and function(*18, 23, 24*) of nucleic acids is well documented in contemporary literature. Note that, such nucleobase protonation remains ‘invisible’ even in the high resolution X-ray crystal structures, and can only be inferred through circumstantial evidences. Our QM studies(*25*) suggest that, the charge redistribution caused by N7 protonation in guanine is qualitatively equivalent to that caused by the coordination of N7 with Mg^2+^ ion.(*13*) It is therefore hypothesized that, guanine N7 protonation may act as a stabilizing factor for those instances of G:C W:W Trans pair, which do not show posttranscriptional modification or metal ion coordination at guanine’s hoogsteen edge or do not take part in higher order interactions.(*17*)

The importance of Mg^2+^ in tRNA folding has been known long since,(*26–28*) and recent studies(29, 30) have also highlighted its role in stabilizing essentially all large RNAs including transition states of some ribozymes.(*31, 32*) Several recent theoretical studies have focused on understanding the Mg^2+^ binding architecture and its exact role in RNA’s structure and function.(*33–37*) However, proper care should be exercised while drawing conclusions based on these studies, as the difference in electron density maps alone can not distinguish between Na^+^, H_2_O and Mg^2+^.(*33*) For example, a very recent article by Leonarski *et al.* has suggested that Mg^2+^ ions assigned near imino nitrogens are often suspect.(*38*) Their results in conjunction with an earlier report by Zheng *et al.*(*33*), on magnesium-binding architectures in RNA crystal structures, suggest that Mg^2+^ coordination with nucleobases takes place via different binding modes in RNA. At the same time, among all the nucleobase atoms, the propensity of Mg^2+^ binding is significantly high for O6 of guanine.(*33, 38*) Further, water molecules in the first coordination shell of Mg^2+^ are highly polarized and have specific roles in mediating Mg^2+^–RNA interactions.(*33, 39*) We have already discussed that, charge redistribution in guanine caused by protonation at N7 can stabilize the RWC geometry of G:C W:W Trans pair. In principle, interaction of the highly acidic first shell water molecules^3^ with N7 of guanine should cause similar charge redistribution and therefore can act as a stabilizing factor for the RWC geometry.

It may be noted that, only Mg^2+^ binding to the N7 of guanine has been studied in the context of the G:C W:W Trans geometry. Therefore, it is necessary to investigate the effect of other binding modes of Mg^2+^ with the hoogsteen edge of guanine and understand their influence on G:C W:W Trans base pairing. In this work, we have analyzed high resolution crystal structures of RNA to identify instances of G:C W:W Trans pair, where Mg^2+^ interacts with the hoogsteen edge of guanine via different binding modes, such as, (a) direct binding at N7, (b) direct binding at O6, (c) simultaneous binding at both N7 and O6 and (d) interaction via hydrogen bonding with the water molecules of the first coordination shell. We also have presented a molecular level understanding of the chemistry of such interactions and their impact on the geometry and stability of G:C W:W Trans base pairs on the basis of DFT calculations.

## Methods

We have curated all the RNA X-ray crystal structures (total 1873 structures) which have a resolution less than 3.5 Å and available in the PDB server on September, 2015. We have also curated all the solution NMR structures containing RNA fragments (total 591 structures) and available in the PDB server on September, 2015. We will refer to this set of 2464 structures as ‘large dataset’ in the following text. To have an unbiased statistics of structural features, we have also considered two non-redundant sets of RNA crystal structures. One is provided by the Nucleic Acid Database (NDB)(*41*) and the other is provided by the HD-RNAS(*42*). We have applied a resolution cut-off of 3.5 Å and shortlisted 838 structures available in the version 1.89 of the NDB database. This set of 838 structures will be referred as ‘NDB dataset’ in the following text. The representative structures present in HD-RNAS are decided upon after taking into account length, R-factor, resolution and sequence similarity. We have further shortlisted 167 RNA crystal structures after applying a resolution cut-off of 3.5 Å and length cut-off of 30 nucleotides in order to exclude the small synthetic RNA constructs. We will refer to this set of 167 structures as ‘HD-RNAS dataset’ in the following text. All the PDB Ids of the crystal structures of the corresponding data sets are listed in the Supporting Information.

For a given RNA crystal structure, first we have shortlisted all the guanine residues which have one or more Mg^2+^ ions within 6.5 Å of their respective hoogsteen edge carbonyl oxygen (O6) and/or imino nitrogen (N7) atoms. These short listed Mg^2+^—nucleobase pairs may further be categorized into two classes depending on the mode of interaction between Mg^2+^ and nucleobase atoms (O6/N7) — (i) direct interaction, and (ii) water mediated interaction. We have taken the ideal bond length of Mg–O and Mg–N bonds as 2.08 Å and 2.20 Å, respectively. These are the mean distances observed in the Cambridge Structural Database(*43*) and also considered in earlier literature.(*33*) With a buffer length of 0.5 Å, if the Mg–O6 distance is found to be ≤ 2.70 Å and/or the Mg–N7 distance is found to be ≤ 2.58 Å, we have considered the corresponding Mg^2+^—nucleobase interaction as a direct interaction. On the other hand, we have considered it as a water mediated interaction, if a water molecule (O*_W_*) is found between the Mg^2+^ and the nucleobase atom in such a way that (i) it is directly bound to Mg^2+^ (i.e., Mg^2+^−O*_W_* distance ≤ 2.70 Å) and (ii) there is a possibility of hydrogen bonding interaction between the water molecule and nucleobase atom i.e., O*_W_*–O6/N7 distance ≤ 3.5 Å, the well accepted cut-off for hydrogen bond donor-acceptor distance for biological systems.(*44, 45*) This algorithm is pictorially depicted in the Figure S1 of Supporting Information.

Next, we have analyzed the given RNA crystal structure with the BPFind software(*46*) and identified whether the Mg^2+^ bound nucleobases are part of any G:C W:W Trans type base pairing interaction or not. BPFind is a well accepted(*47*) precursor atom based algorithm, which identifies two nucleobases as a base pair if there are at least two conventional hydrogen bonds (N-H…N, N-H…O, O-H…N, O-H…O, C-HH…N and C-HH…O type) present between them. We have applied the following cut-offs to detect only ‘good’ base pairing interactions – (i) cutoff distance of 3.8 Å between the acceptor and donor atoms, (ii) cutoff angle of 120.0° for checking planarity of precursor atoms and linearity of the hydrogen bonds and (iii) cutoff ‘E-value’ of 1.8 to signify the overall distortion and maintain a good base pairing geometry.

We have extracted the coordinates of nucleobases and Mg atoms from the crystal structures and modeled the Mg^2+^ coordinated G:C W:W Trans base pairs by adding hydrogen atoms and waters (at the first coordination shell of Mg^2+^) manually using GaussView(*48*) software. As per common practice,(*49*) we have substituted the sugar moiety by methyl group to reduce the computational cost without compromising with the accuracy of results. All the base pairs and the corresponding monomers are geometry optimized at six different DFT functional. The first one is B3LYP,(*50–52*) as it is arguably the most popular hydrid GGA functional (20% HF exchange) implemented in studying base pairing systems.(*15, 53*) Hessian calculations were performed for all the optimized geometries to confirm that they are not associated with any imaginary frequencies.

Interaction energy of these systems (at DFT level) has been calculated as, 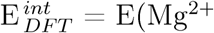 bound base pair) - E(Mg^2+^ bound guanine) - E(cytosine) + BSSE, where E(X) represents the electronic energy corresponding to the optimized geometry of isolated X and BSSE represents the basis set superposition error calculated using counterpoise method.(*54*) All the DFT calculations have been performed using 6-311G++(d,p) basis set for Mg and 6-31++g(2d,2p) basis set for other atoms.(*55*) For the B3LYP optimized geometries we have also calculated the BSSE corrected interaction energy at MP2 level and denoted it as E^*int*^. Further, we have studied the solvent screening of the electrostatic component of the interaction energy by using the conductor-like polarizable continuum model (CPCM),(*56, 57*) with water as solvent (*ε* = 78.4) and denoted it 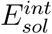. as Note that, CPCM uses united atom topological model to define the atomic radii and is found to be appropriate for polar liquids.(*58*) All MP2 calculations have been performed using aug-cc-pVTZ basis set for Mg and aug-CC-pVDZ basis set for other atoms.(*59*) Kitaura-Morokuma decomposition analysis(*60*) were performed (using GAMESS-US package(*61*)) to study the partitioning of the two body intermolecular interaction energies into electrostatic, polarization, charge transfer and higher order coupling terms, within the Hartree-Fock approximation. To study the charge distribution of a system, we have performed Natural Population Analysis(*62, 63*) using the NBO package(*64*) implemented in Gaussian 09 package. All other QM calculations are also done using Gaussian 09 package.(*65*)

NUPARM package(*66,67*) has been used to characterize the geometry of optimized base pairs on the basis of three translational (shear, stretch and stagger) and three rotational (buckle, propeller twist and opening) parameters. Structural alignment was performed using VMD,(*68*) We have developed a Python based program to detect Mg^2+^ coordinated base pairs in RNA crystal structures.

## Results and Discussions

### Context analysis of G:C W:W Trans base pairs in RNA crystal structures

Two nonredundent datasets of RNA crystal structures analyzed in this work, HD-RNAS and NDB contain 116 and 86 instances of G:C W:W Trans, respectively. However, 2101 instances of G:C W:W Trans have been detected in the large set of RNA crystal structures. We have analyzed the context of occurrence of these instances and found that G:C W:W Trans occurs in 14 different contexts (Table 1), Among these 14 contexts, 8 show Mg^2+^ coordination at the hoogsteen edge of guanine (Table 1), Apart from the well studied Levitt pair (context 1), Mg^2+^ coordinated G:C W:W Trans is observed in 23S rRNA of *Escherichia coli* (context 2d and context 10), 23S rRNA of *Thermus thermophilus* (context 4b, context 5), 23S rRNA of *Haloarcula marismortui* (context 3c), 23S rRNA (context 2e) and 5.8S rRNA (context 13) of *Saccharomyces cerevisiae*. A detailed picture of the occurrence contexts are given in the Supporting Information (Figure S2), which shows that Mg^2+^ coordinated G:C W:W Trans pairs are involved in various structural motifs, such as kissing loop, internal loop, junction loop and also mediate loop-loop interactions.

**Table 1:** Context of occurrence of G:C W:W Trans base pair in the large set of RNA crystal structures. Average (*E̅*) and standard deviation (SD) of the E-value (calculated by BPFIND software) have been reported for each context.

| | RNA type | Species | Base pair | Mg<sup>2+</sup><br>coordination | Occ.<br>Freq. | E value<br>$\bar{E}$ SD | | Context of occurrence |
| --- | --- | --- | --- | --- | --- | --- | --- | --- |
| 1. | tRNA | various species | 15G:48C | yes | 396 | 0.67 | 0.34 | Levitt base pair |
| 2. | (a) 23S rRNA | *D. radiodurans* | 2508G:2454C |  | 15 | 1.01 | 0.55 | Kissing loop in Domain V |
|  | (b) 23S rRNA | *T. thermophilus* | 2541G:2487C |  | 19 | 0.64 | 0.24 |  |
|  | (c) 23S rRNA | *H. marismortui* | 2564G:2510C |  | 65 | 0.90 | 0.27 |  |
|  | (d) 23S rRNA | *E. coli* | 2529G:2475C | yes | 177 | 0.77 | 0.32 |  |
|  | (e) 25S rRNA | *S. cerevisiae* | 2898G:2844C | yes | 34 | 0.55 | 0.15 |  |
| 3. | (a) 23S rRNA | *D. radiodurans* | 1809G:1791C |  | 7 | 0.89 | 0.21 | Internal Loop in Domain IV |
|  | (b) 23S rRNA | *T. thermophilus* | 1817G:1800C |  | 209 | 0.80 | 0.17 |  |
|  | (c) 23S rRNA | *H. marismortui* | 1873G:1856C | yes | 65 | 0.54 | 0.10 |  |
| 4. | (a) 23S rRNA | *D. radiodurans* | 2484G:2589C |  | 7 | 0.86 | 0.28 | 5-way junction loop in Domain V |
|  | (b) 23S rRNA | *T. thermophilus* | (a) 2505G:2610C<br>(b) 2517G:2622C | yes | 199<br>22 | 0.73<br>0.62 | 0.25<br>0.13 | 5-way junction loop connecting Domain 0 and Domain V |
| 5. | 23S rRNA | *T. thermophilus* | 1271G:1615C | yes | 212 | 0.62 | 0.20 | Connecting a junction loop (gua) and a hairpin loop (cyt) in Domain III |
| 6. | 23S rRNA | *T. thermophilus* | (a) 1929G:1925C<br>(b) 1951G:1947C |  | 205 | 0.50 | 0.20 | 4-way junction loop in Domain IV |
| 7. | 23S rRNA | *T. thermophilus* | 430G:234C |  | 173 | 0.45 | 0.14 | Connecting two junction loops in Domain I |
| 8. | (a) 23S rRNA | *H. marismortui* | 1190G:1186C |  | 60 | 0.83 | 0.26 | 3-way junction loop in Domain II |
|  | (b) 25S rRNA | *S. cerevisiae* | 1261G:1257C |  | 34 | 0.58 | 0.36 |  |
| 9. | 23S rRNA | *H. marismortui* | 1683G:1377C |  | 65 | 0.36 | 0.09 | Connect 2 junction loops within Domain III |
| 10. | 23S rRNA | *E. coli* | 1360G:2214C | yes | 53 | 0.92 | 0.33 | kissing loop interaction connecting Domain III and Domain V |
| 11. | (a) 23S rRNA | *D. radiodurans* | 1284G:1631C |  | 20 | 0.92 | 0.34 | Helix-helix interaction between Domain III (gua) and Domain 0 (cyt) |
|  | (b) 23S rRNA | *T. thermophilus* | 1318G:1662C |  | 20 | 0.56 | 0.11 |  |
| 12. | 23S rRNA | *T. thermophilus* | 1847G:1830C |  | 20 | 0.69 | 0.11 | Bulge-Helix interaction in Domain IV |
| 13. | 5.8S rRNA | *S. cerevisiae* | 39G:106C | yes | 17 | 1.00 | 0.11 | Loop-loop interaction (a part of bifurcated triple) |
| 14. | 18S rRNA | *S. cerevisiae* | 720G:717C |  | 7 | 1.60 | 0.20 | Hairpin loop in Domain C |

Analysis of the large set of RNA crystal structures shows that 36.2% of the total instances of G:C W:W Trans pair have at least one Mg^2+^ within 6.5Å of either N7 or O6. This abundance of Mg^2+^ near G:C W:W Trans is remarkable as the ratio is even higher than the hoogsteen edge of canonical G:C W:W Cis pairs (32.2%, with Mg^2+^ near guanine). Interestingly, distribution of occurrence frequency of G:C W:W Trans over the Mg-N7 (Figure 2A) and Mg-O6 distances (Figure 2B) are distinctly different. Occurrence frequency of G:C W:W Trans pairs that have a Mg^2+^ within 6.5Å of N7 (698) is significantly greater than the occurrence frequency of G:C W:W Trans pairs that have a Mg^2+^ within 6.5Å of O6 (518). However, for the latter case, the propensity of occurrences of Mg^2+^ within 3.0Å of the base atom (i.e., close to the Mg-O/Mg-N covalent bond length) is remarkably high. This indicates towards the possibility of direct coordination of guanine’s O6 with Mg^2+^. In contrast, frequency distribution over Mg-N7 distance has a high population at longer distances (~4.5Å and ~5.7Å). This indicates towards the possibility of Mg^2+^ binding at N7 via hydrogen bonding interactions with the water molecules of the first coordination shell. Implication of these new Mg^2+^ binding modes have not been studied earlier in the context of G:C W:W Trans pair. We have, therefore, analyzed the RNA crystal structures to identify these new modes of Mg^2+^ binding with guanine’s hoogsteen edge in G:C W:W Trans pairs.

**Figure 2:**
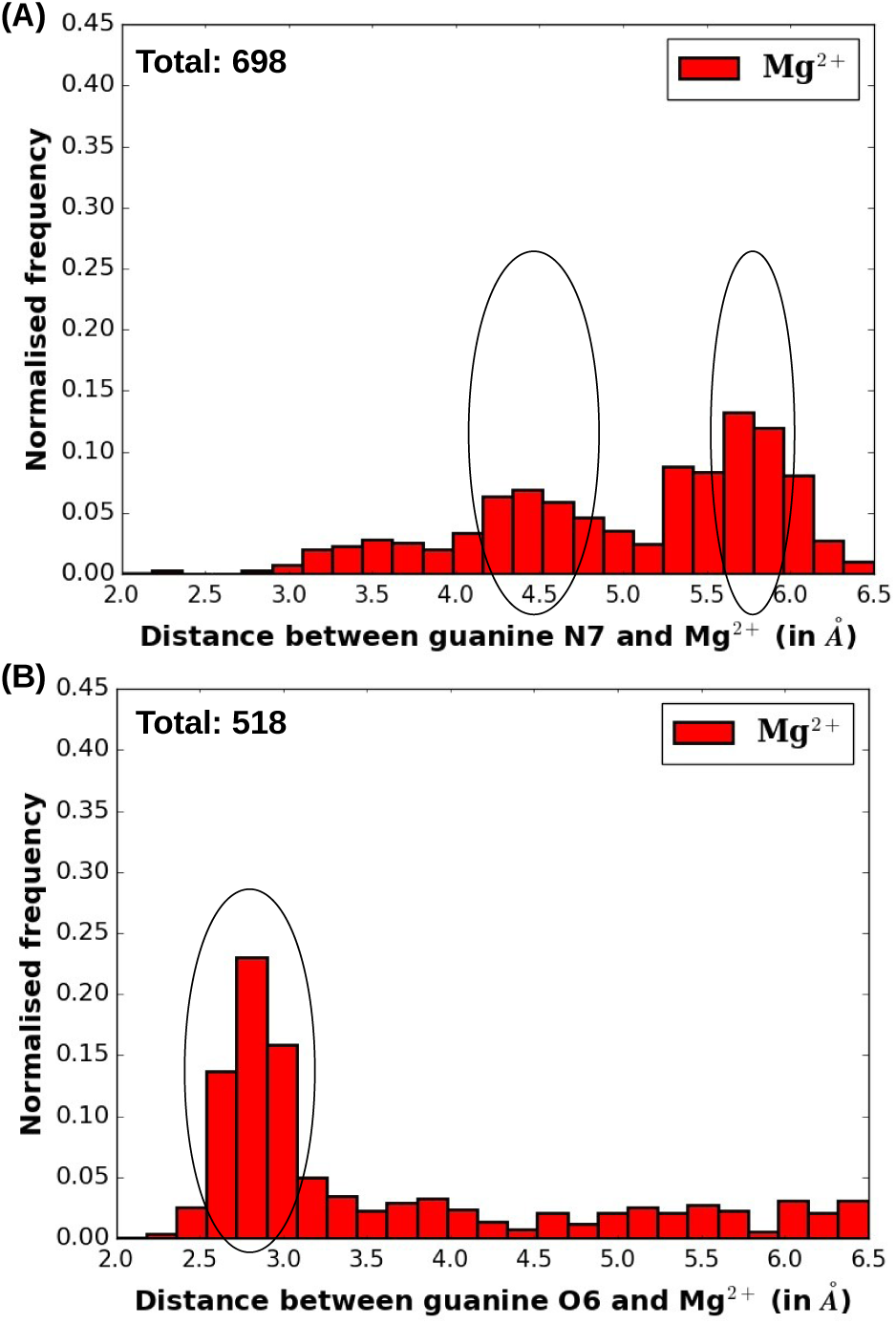
Distribution of occurrence frequency (normalized) of G:C W:W Trans pair that have atleast one Mg^2+^ within 6.5Å of the nucleobase atom (N7/O6) with respect to (A) N7-Mg^2+^ distance and (B) O6-Mg^2+^ distance.

Based on inter-atomic distances calculated from crystal coordinates, it is not possible to identify the exact binding mode for all the cases. In the large set of RNA crystal structures, we have identified only 1 instance of direct Mg^2+^ binding at N7, 23 instances of direct Mg^2+^ binding at O6 and 86 instances of Mg^2+^ binding via the water molecules of first coordination shell. We have characterized the geometry of these Mg^2+^ bound G:C W:W Trans pairs by the three translational (shear, stretch and stagger) and three rotational (buckle, propeller twist and opening) parameters as shown in Figure 3A, Distribution of these instances in the Buckle-Propeller Twist space (Figure 3B) shows that, water mediated Mg^2+^ binding at the hoogsteen edge of guanine (red circle) and direct Mg^2+^ binding at O6 (blue circle) are usually associated with a nonplanar geometry characterized by high buckle and/or propeller twist values. On the other hand, direct Mg^2+^ binding at N7 is associated with a relatively planar geometry (cyan circle). Distribution in the Buckle-Open (Figure 3C) and Open-Propeller Twist (Figure 3D) spaces shows no significant variation in the opening angle. However, data points corresponding to the water mediated Mg^2+^ binding (red circle) and direct Mg^2+^ binding at O6 (blue circle) are distinctly clustered at the high buckle and propeller twist values (both positive and negative) in Figure 3C and 3D, respectively. The planar geometry of G:C W:W Trans corresponding to direct Mg^2+^ binding at N7 is consistent with the optimized geometries reported in earlier QM based studies, (*13, 14*) Can the nonplanar geometries corresponding to the new Mg^2+^ binding modes also be explained on the basis of their ground state electronic structures?

**Figure 3:**
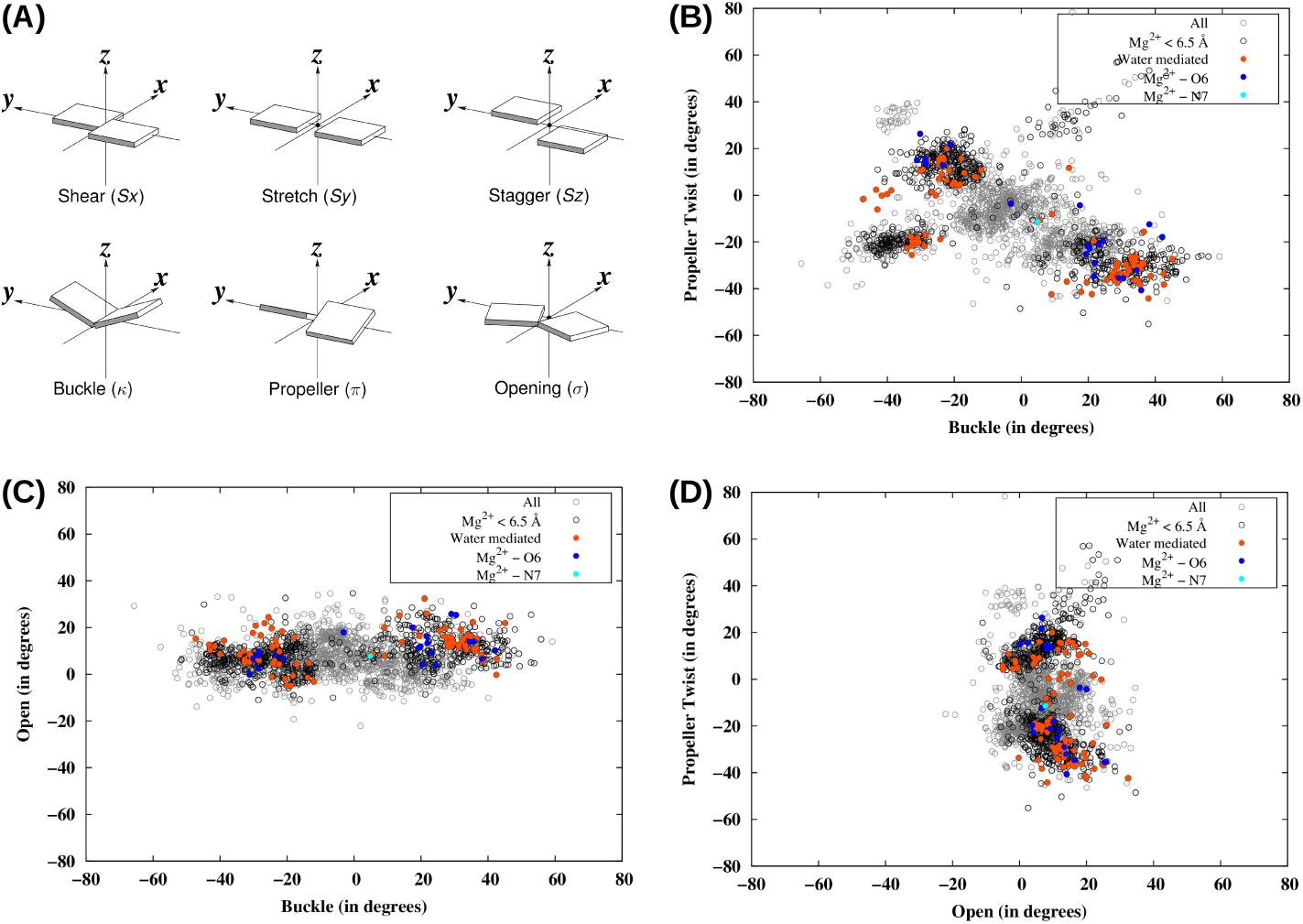
(A) Definition of the three translational (shear, stretch and stagger) and three rotational (buckle, propeller twist and opening) that defines the mutual orientation of the two bases in a base pair (Image source: http://x3dna.org/). Distribution of all instances of G:C W:W Trans base pair detected in the complete set of RNA crystal structures are shown in the (B) Buckle-Propeller Twist, (C) Buckle-Open and (D) Open-Propeller Twist spaces. Filled circles represent the instances for which the exact binding mode of Mg^2+^ (*cyan* - direct coordination at N7, *blue* - direct coordination at O6, *red* - coordination via first shell water molecules) can be identified in the crystal structures. Black empty circles represent instances where Mg^2+^ is present within 6.5Å of N7/O6, but the exact binding mode can not be identified. All other instances are shown in gray empty circles.

### Influence of Mg^2+^ coordination on the geometry and charge distribution of guanine

Apart from the three different Mg^2+^ binding modes observed in crystal structures (e.g., (i) direct Mg^2+^ binding at N7, (ii) direct Mg^2+^ binding at O6 and (iii) Mg^2+^ binding via first shell water molecules), we have modeled one more mode of interaction — (iv) direct binding of Mg^2+^ with both N7 and O6 simultaneously. Optimized geometry (at B3LYP) of Mg^2+^ bound guanine corresponding to these four different binding modes are shown in Figure 4, Comparison with the optimized geometry of normal guanine (Figure 4A) reveals that, the major change in the geometry takes place at the exocyclic amino group. In all the four cases (Figure 4C-F), the change is characterized by depyramidalization of the amino group and shortening of the C2-N2 bond, which is indicative of a sp^3^ → sp^2^ transition of N2’s hybridization state. This can be explained by the conjugation of the lone pair of the exocyclic amino group with the ring *π* system (Figure 4G), As discussed earlier for guanine N7 protonation,(*17*) positive charge build up at the hoogsteen edge of guanine favors such charge delocalization by stabilizing the resonance canonical structure labeled as (ii) in Figure 4G, Such Mg^2+^ binding induced charge delocalization is also reflected in the reduced HOMO-LUMO gaps (ΔE_*H–L*_) in Mg^2+^ bound systems, e.g., at B3LYP level ΔE_*H–L*_(normal guanine) = 5.1eV, ΔE_*H–L*_(direct Mg^2+^ binding at N7/O6/both) = 4.9eV, ΔE_*H–L*_(water mediated Mg^2+^ binding) = 4.8eV.

**Figure 4:**
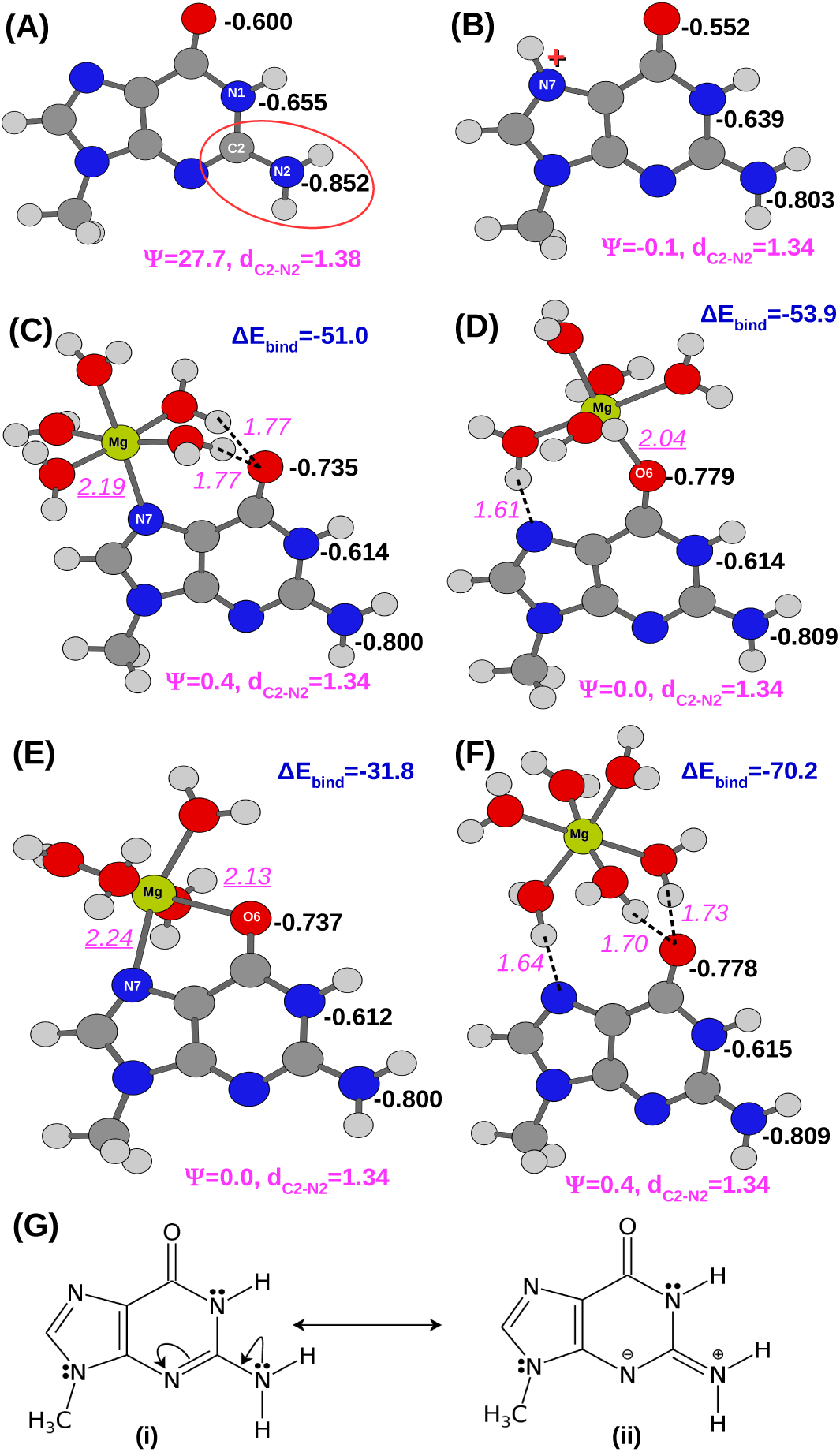
B3LYP optimized geometries of (A) normal guanine, (B) N7 protonated guanine, (C) guanine with direct Mg^2+^ coordination at N7, (D) guanine with direct Mg^2+^ coordination at O6, (E) guanine with direct Mg^2+^ coordination at both N7 and O6, and (F) guanine with Mg^2+^ coordinating at the hoogsteen edge via the water molecules of its first coordination shell. The improper dihedral angle *ψ* measures the extent of pyramidalization of the amino group (see Figure S3 of Supporting Information). d_*C*2–*N*2_ represents the C2-N2 bond length. Hydrogen bonds are shown as broken lines. All inter atomic distances are reported in Å in magenta text. Mg-N7/O6 distances are underlined. Partial charges obtained from natural population analysis are reported (in a.u) for the hydrogen bond donor and acceptor atoms of the Watson-Crick edge (in black text). ΔE_*bind*_ represents the binding energy (in kcal/mol, MP2) between the hydrated Mg^2+^ ion and guanine. (G) Conjugation of the lone pair of the exocyclic amino group with the ring *π* system is shown.

Mg^2+^ induced charge redistribution modulates the hydrogen bonding potential of the hydrogen bond donor and acceptor sites of the Watson-Crick edge. On the basis of NBO charges (q) shown in Figure 4, it is evident that both N7 protonation and Mg^2+^ coordination reduces the electron density over the hydrogen bond donor sites (N1 and N2) and thus improves their hydrogen bonding potential. To quantify this variation let us introduce the parameter, Δq = q(Mg^2+^ coordinate guanine) - q(normal guanine). Δq corresponding to N1 and N2 respectively, are similar for all the four Mg^2+^ binding modes. However, the change in the hydrogen bond acceptor site (O6) is opposite for N7 protonation and Mg^2+^ interaction. Across all the four binding modes studied, Mg^2+^ interaction increases the electron density over O6, making it a stronger hydrogen bond acceptor. Interestingly, the four binding modes can be classified into two groups on the basis of their corresponding Δq values, — (*Group I*) direct binding at N7 and at both N7 and O6 Δq = 0.135a.u. and 0.137a.u. for Figure 4C and 4E, respectively) and, (*Group II*) direct binding at O6 and water mediated binding Δq = 0.179a.u. and 0.178a.u. for Figure 4D and 4F, respectively). The same grouping is also valid for the extent of decrease in electron density over the hydrogen atoms of the Watson-Crick edge (Table S1 of Supporting Information). It is noteworthy that, earlier works have considered the decrease in electron density over the amino hydrogens as the key factor responsible for the stabilization of the RWC geometry on Mg^2+^ coordination, since it reduces the electrostatic repulsion between amino groups of guanine and cytosine.(13) Therefore, on the basis of Mg^2+^ induced charge redistribution we can expect that, the influence of Mg^2+^ coordination on the G:C W:W Trans base pairing will be different for the two sets of conditions mentioned above, *Group I* and *Group II*.

An important observation is that, stability of the hydrated Mg^2+^-guanine complex is highly dependent on the binding mode (Figure 4C-F), Water mediated Mg^2+^ interaction is significantly stronger (E*_bind_* = -70.2 kcal/mol) than any direct Mg^2+^ interaction mode. Due to restriction in the formation of the octahedral geometry, direct Mg^2+^ binding at both N7 and O6 (E*_bind_* = -31.8 kcal/mol) is significantly weaker than for direct Mg^2+^ binding at N7 (E_*bind*_ = -51.0 kcal/mol) and O6 (E_*bind*_ = -53.9 kcal/mol). Here, E_*bind*_ for a (Mg^2+^, nH_2_O) bound guanine has been calculated as, E_*bind*_ = E(guanine bound to Mg^2+^,nH_2_O) + (6-n)E(H_2_O) - E(Mg^2+^,6H_2_O), where E(X) corresponds to the ground state electronic energy of the species X.

### Influence of different Mg^2+^ coordination modes on the geometry and stability of G:C W:W Trans pair

From the B3LYP level optimized geometries shown in Figure 5, it is clear that, RWC geometry of G:C W:W Trans pair is not stable in gas phase and converges to the bifurcated geometry on energy optimization (Figure 5A), Positive charge build up at the hoogsteen edge of guanine in the form of direct N7 protonation (Figure 5B), direct Mg^2+^ binding (Figure 5C-E), or, water mediated Mg^2+^ interaction (Figure 5F) stabilizes the RWC geometry. Figure 5 also shows the BSSE corrected interaction energy calculated at MP2 level for the B3LYP optimized geometries. In general, gas phase interaction energy of these positively charged systems are dominated by electrostatic component (HF component of 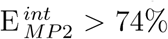 for all the systems, Table S2 in Supporting Information). We have, therefore, incorporated the effect of solvent screening on the interaction energy (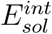) of the gas phase optimized geometries. 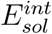 values shown in Figure 5 suggest that, the RWC geometry stabilized by positive charge build up at hoogsteen edge is 3.3 to 6.6 kcal/mol stabler than for the bifurcated geometry. Among the 3 direct Mg^2+^ binding modes, direct binding at O6 results in the strongest pair (E^*int*^ = -43.0kcal/mol, 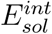 = -16.1kcal/mol), probably due to the extra hydrogen bond between one first shell water and O2 of cytosine. However, interaction energy of G:C W:W Trans with water mediated Mg^2+^ binding is lower than all three direct Mg^2+^ coordinated models. This relative order of stability of the G:C W:W trans pairs corresponding to the four Mg^2+^ binding modes is consistent with the interaction energies calculated at six different DFT functionals (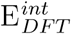 in Table **??**).

**Figure 5:**
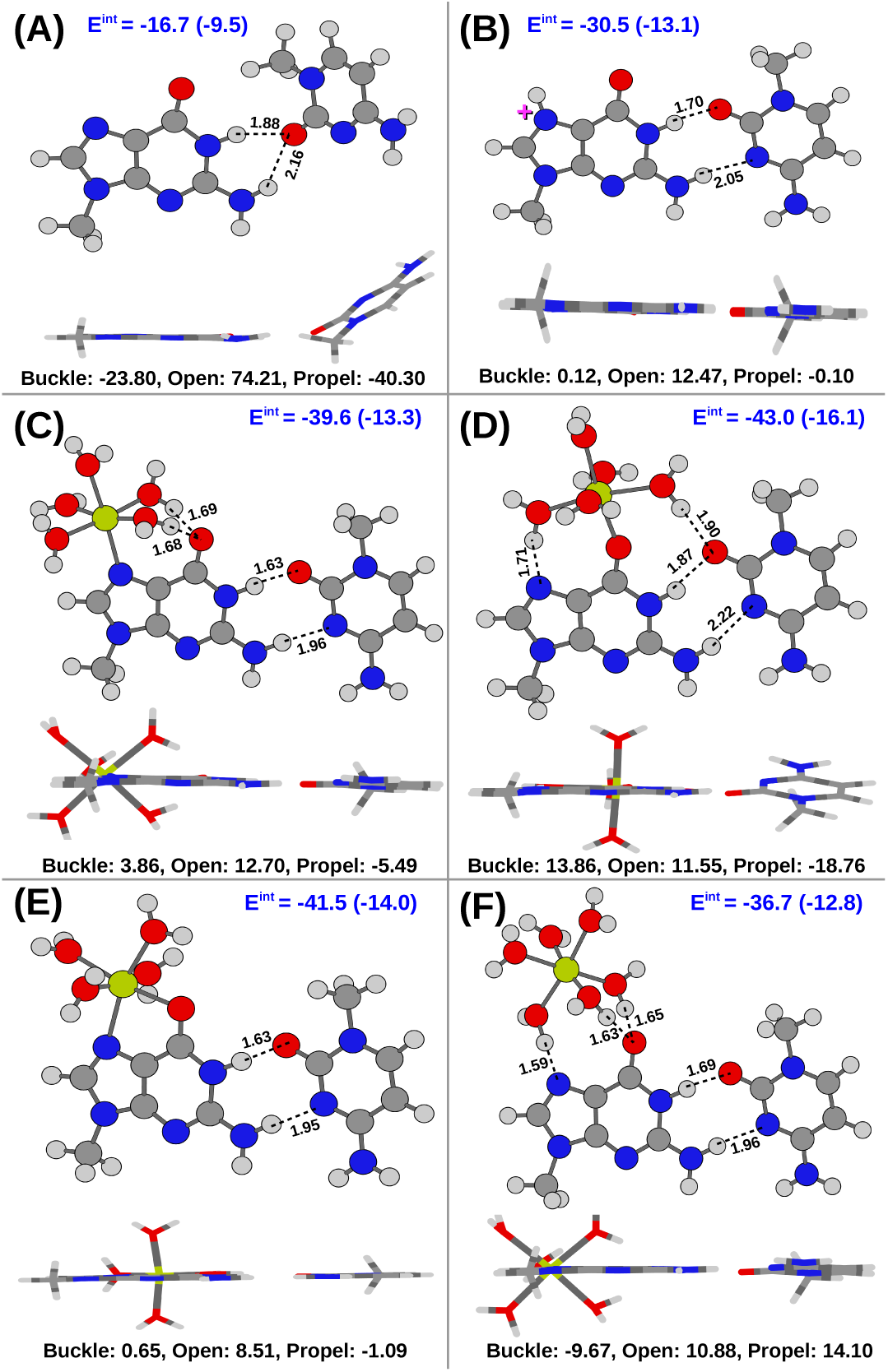
B3LYP optimized geometries of G:C W:W Trans pair with (A) normal guanine, (B) N7 protonated guanine, and (C – F) Mg^2+^ coordinated guanine are shown. Different modes of Mg^2+^ coordination are, direct Mg^2+^ coordination at (C) only N7, (D) only O6, (E) both N7 and O6, and (F) coordination via the water molecules of first coordination shell. For each case, a lateral view of the optimized structure is also shown to highlight the extent of nonplanarity of the interaction. The same has been quantified by Buckle, Open and Propeller Twist parameters. Hydrogen bonds are represented by broken lines along with the distances between the hydrogen and and the acceptor atom in Å. BSSE corrected interaction energies (E^*int*^) calculated at MP2 level have been reported in kcal/mol. E^*int*^ values calculated in solvent phase are given in parenthesis.

Interestingly, as expected from the Mg^2+^ coordination induced charge redistribution in guanine, the optimized geometry of G:C W:W Trans with *Group I* type Mg^2+^ interaction is significantly different from that with *Group II* type Mg^2+^ interaction. For the latter group, the optimized geometry is nonplanar having high buckle and propeller twist (Figure 5D, 5F), whereas the former results in a planar geometry (Figure 5C, 5E) similar to guanine N7 protonation (Figure 5B). The comparison is clearly depicted in Figures 6A and 6B, where the optimized geometries are structurally aligned with respect to the guanine residue. Optimized geometries corresponding to direct Mg^2+^ binding at O6 (green) and Mg^2+^ binding via first shell water molecules (red) bend out of the plane, whereas, optimized geometries corresponding to N7 protonation (blue), direct Mg^2+^ coordination at N7 (magenta), and at both N7 and O6 (yellow) remain planar.

**Figure 6:**
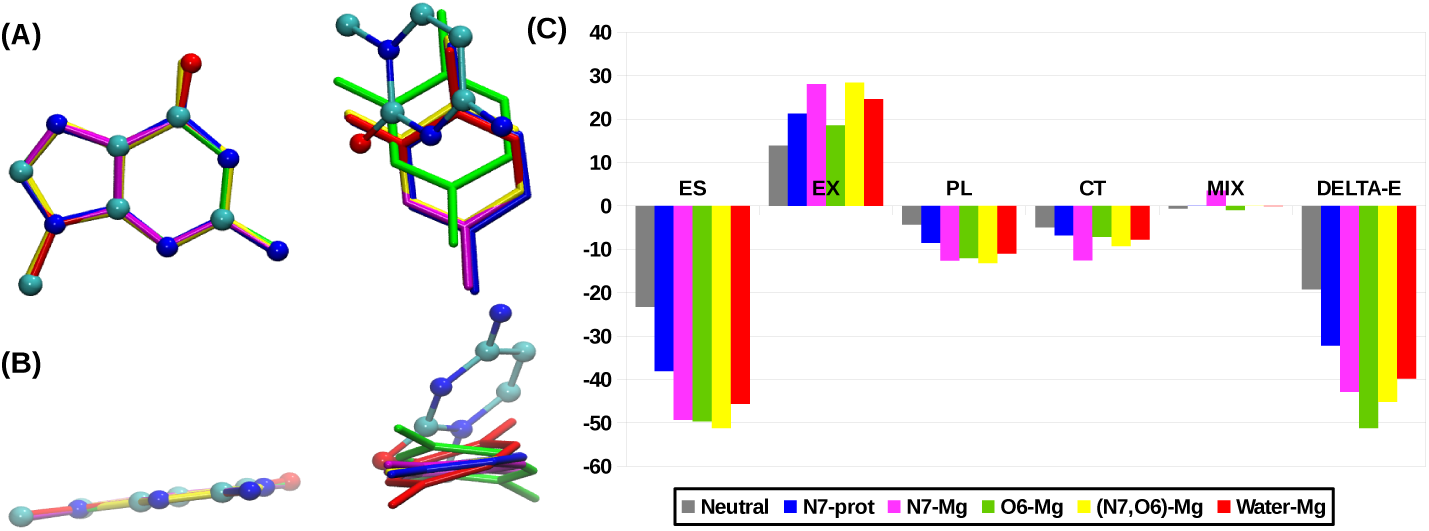
(A) B3LYP optimized geometries of G:C W:W Trans pair with normal (ball & stick), N7 protonated (blue) and Mg^2+^ bound guanine are superposed with respect to the guanine residue. (B) Same structural superposition is shown from a lateral view. Planar geometry for Mg^2+^ coordination at only N7 (magenta) and at both N7 and O6 (yellow) and, nonplanar geometry for Mg^2+^ coordination at only O6 (green) and water mediated Mg^2+^ coordination (red) are noticed. (C) Different components (ES = electrostatic, EX = exchange repulsion, PL = polarization, CT = charge transfer, MIX = higher order coupling) of the total interaction energy (DELTA-E) are shown.

### Nonplanarity of G:C W:W Trans on direct Mg^2+^ binding at O6 and Mg^2+^ binding via first shell waters

Geometry optimization result in a buckled and propeller twisted geometry for ( *case 1*) direct Mg^2+^ binding at O6 and (*case 2*) Mg^2+^ binding via first coordination shell water molecules (Table **??**). Inter base hydrogen bonding distances corresponding to *case 1* and *case 2* are longer than that for the N7 protonated case. This indicates that, the buckled and propeller twisted geometry moves the bases away from each other. Again, for (*case 3*) direct Mg^2+^ coordination at N7 (Figure 5C) and (*case 4*) at both N7 and O6 (Figure 5E), the optimized geometries remain planar. The inter base hydrogen bonding distances corresponding to *case 3* and *case 4* are shorter than that for N7 protonated case. This indicates that, the planar geometry brings the bases close to each other.

Decomposition of the interaction energies (Figure 6C) of the B3LYP optimized geometries show that, in comparison to N7 protonation (blue bar), Mg^2+^ coordination significantly increases the electrostatic component for all the four cases. However, the corresponding charge transfer components are relatively higher for *case 3* (magenta bar) and *case 4* (yellow bar). As a result, although the planar geometry of *case 3* and *case 4* bring the bases closer to each other, the subsequent increase in the exchange repulsion component gets compensated by the charge transfer component. On the other hand, the charge transfer component is relatively lower in *case 1* (green bar) and *case 2* (red bar). Therefore, these two systems tend to attain optimum stability by adopting a buckled and propeller twisted geometry, which results in the movement of the two bases away from each other, thus avoiding any consequent increase of the exchange repulsion component.

### Consistency with the crystal structure

Optimized geometries corresponding to different Mg^2+^ binding modes are consistent with the trends observed in RNA crystal structures. As discussed in the previous section, the identified G:C W:W Trans pairs with direct Mg^2+^ binding at O6 and water mediated Mg^2+^ binding is also associated with a buckled and propeller twisted geometry in the crystal structure. Note that, in the crystal structures, apart from the mode of Mg^2+^ coordination, there are several other factors that determine the planarity of a base pairing interaction. However, in this particular case, the consistency between the crystal geometry and the optimized geometry is remarkable. It shows that Mg^2+^ can fine tune the geometry of base pairing interactions. The small difference between the interaction energy of G:C W:W Trans with (a) direct Mg^2+^ binding at N7 and (b) Mg^2+^ binding via first shell waters (Δ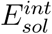 = 0.5keal/mol) further suggests that, such Mg^2+^ binding based geometric modulation can take place without changing the stability of the overall system.

## Conclusions

G:C W:W Trans base pairs occur frequently in 23S rRNA of all the four species available in our dataset. Occurrence of G:C W:W Trans is also observed in 5.8S rRNA of Saccharomyces cerevisiae and in tRNAs (the popular Levitt base pair) of various species. All these occurrences can be classified into 14 contexts. Among them, Mg^2+^ binding at the hoogsteen edge of guanine is observed for 8 contexts, which are integrally associated with a variety of structural motifs including different types of loops and their interactions. Sensitivity of the geometry of these base pairs to the binding mode of Mg^2+^, as described in this work, may imply significant consequences of binding mode variation on the functional role of those structural motifs. Such sensitivity towards Mg^2+^ binding modes need not be restricted to G:C W:W Trans pairs alone. In fact binding of magnesium or other divalent ions to other noncanonical base pairs may also result in similar structural variations. Our results thus open up an additional avenue for researching the structural complexity and functional diversity of RNA molecules.

## Acknowledgement

A.H. acknowledges CSIR for SRF support. A.M. and A.H. thank DBT, Government of India project BT/PR-14715/PBD/16/903/2010 for partial funding and financial support. A.M. and D.B. thank DBT, Government of India project BT/PR-11429/BID/07/271/2008 for supporting computational infrastructure.

## Supporting Information Available

The following files are available free of charge.

1. GCWWT-Mg-Supporting-Information.pdf: Pictorial description of the algorithm that has been implemented to detect different Mg^2+^ bound G:C W:W Trans pairs, details of the context of occurrence of Mg^2+^ bound G:C W:W Trans pairs, definition of the angle *ψ*, partial charges over the hydrogen atoms of the WC edge of guanine, details of interaction energies, and name of the PDB files present in the 3 data sets studied.
2. Optimized-Geometries-B3LYP.tar.gz: Cartesian coordinates of the optimized geometries of the base pairs and their monomers in .pdb format.

## Footnotes

1 This bifurcated geometry can be classified as G-C Ww/Bs *trans*, according to the Leontis and Westhof nomenclature (LW) extended for bifurcated geometries.(*5, 12*) However, in this work, to annotate the base pairing interactions we have followed a nomenclature which is slightly different from the LW nomenclature. If the edge X of base A interacts with the edge Y of base B in *cis* (or *trans*) orientation, that is annotated as A:B X:Y Cis (or, A:B X:Y Trans). Therefore, the crystal geometry of Levitt base pair is annotated as G:C W:W Trans

2 The pK_*a*1_ values of adenine (~4.1), guanine (~3.2) and cytosine (~4.4) are usually 2-3 units away from neutrality.(*18*)

3 pK_*a*_ of Mg^2+^(H_2_O)6 = 11.4, pK_*a*_ of Na^+^(H_2_O)_6–8_= 14.4, pK_*a*_ of H_2_O_*bulk*_ = 15.7.(*40*)

